# Improved assembly and variant detection of a haploid human genome using single-molecule, high-fidelity long reads

**DOI:** 10.1101/635037

**Authors:** Mitchell R. Vollger, Glennis A. Logsdon, Peter A. Audano, Arvis Sulovari, David Porubsky, Paul Peluso, Aaron M. Wenger, Gregory T. Concepcion, Zev N. Kronenberg, Katherine M. Munson, Carl Baker, Ashley D. Sanders, Diana C.J. Spierings, Peter M. Lansdorp, Urvashi Surti, Michael W. Hunkapiller, Evan E. Eichler

## Abstract

The sequence and assembly of human genomes using long-read sequencing technologies has revolutionized our understanding of structural variation and genome organization. We compared the accuracy, continuity, and gene annotation of genome assemblies generated from either high-fidelity (HiFi) or continuous long-read (CLR) datasets from the same complete hydatidiform mole human genome. We find that the HiFi sequence data assemble an additional 10% of duplicated regions and more accurately represent the structure of tandem repeats, as validated with orthogonal analyses. As a result, an additional 5 Mbp of pericentromeric sequences are recovered in the HiFi assembly, resulting in a 2.5-fold increase in the NG50 within 1 Mbp of the centromere (HiFi 480.6 kbp, CLR 191.5 kbp). Additionally, the HiFi genome assembly was generated in significantly less time with fewer computational resources than the CLR assembly. Although the HiFi assembly has significantly improved continuity and accuracy in many complex regions of the genome, it still falls short of the assembly of centromeric DNA and the largest regions of segmental duplication using existing assemblers. Despite these shortcomings, our results suggest that HiFi may be the most effective stand-alone technology for *de novo* assembly of human genomes.

## Introduction

Recent advances in long-read sequencing technologies, including Pacific Biosciences (PacBio) and Oxford Nanopore Technologies (ONT), have revolutionized the assembly of highly contiguous mammalian genomes (Bickhart et al. 2017; Chaisson et al. 2015; Gordon et al. 2016; Huddleston et al. 2017; Jain et al. 2018; Kronenberg et al. 2018; Low et al. 2019; Seo et al. 2016; Steinberg et al. 2016). For example, individual laboratories can now accurately assemble >90% of mammalian euchromatin in less than 1,000 contigs within a few months. However, the generation of high-quality datasets is costly and requires computational resources unavailable to most researchers. Long-read *de novo* assemblies of human samples typically require 20,000–50,000 CPU hours (Chin et al. 2016; Koren et al. 2017) and terabytes of data storage.

With the recent introduction of high-fidelity (HiFi) sequence data from PacBio and the development of the SMRT Cell 8M, the accessibility of *de novo* assembly using single-molecule, real-time (SMRT) sequencing data has significantly improved. With 28-fold sequence coverage of the Genome in a Bottle (GIAB) Ashkenazim sample HG002, Wenger and colleagues demonstrated that it is possible to create a *de novo* assembly comparable to previous long-read assemblies with half the data and one-tenth the compute power (Wenger et al. 2019). While compute time and throughput have improved, there is little comparison of the HiFi assembly quality of HG002 to a previous continuous long-read (CLR) HG002 genome assembly and limited assessment of the more difficult regions of the genome.

Here, we generate 24-fold sequence coverage and produce a *de novo* assembly of a complete hydatidiform mole human genome (CHM13) with HiFi data. We directly compare it to a previous assembly of CHM13 produced with CLR data (Kronenberg et al. 2018). The accurate assembly of the CHM13 genome is valuable for several reasons. First, due to its single-haplotype nature, it allows for better resolution of highly duplicated sequences, including segmental duplications (SDs) and tandem repeats. This 5-8% portion of the genome represents some of the most challenging regions to resolve. Second, its monoallelic nature permits the detection and unambiguous resolution of structural variants (SVs) that are crucial in disease and evolution. Finally, it allows for complete and absolute deduction of the sequence accuracy of a genome assembly [i.e., quality value (QV)] because there is only one haplotype for comparison. As a result, large-insert BAC clone sequences from the same source material can be expected to align at nearly 100% sequence identity and therefore be used to reliably compute the accuracy of different sequencing platforms and assembly approaches.

## Results

### Whole-genome assembly with HiFi versus CLR reads

To assess the utility of PacBio’s HiFi technology (Wenger et al. 2019) for *de novo* assembly, we set out to compare assemblies of the CHM13 genome using either HiFi (generated on the Sequel II platform) or CLR (generated on the RS II platform) data. To do this, we generated 24-fold HiFi circular consensus sequence (CCS) data from four SMRT Cells 8M. Each SMRT Cell produced, on average, 19.1 Gbp of QV > 20 sequence data (range 14-25 Gbp) with an average consensus read length of 10.95 kbp. The long-read sequence data were of high quality, with 61.2% of the quality-filtered CCS reads having an estimated QV > 30 (**Fig. S1**). The generation of HiFi data using the CCS algorithm took on average 12,500 CPU hours for each SMRT Cell 8M.

Using Canu (Koren et al. 2017) (**Methods**), we generated a *de novo* assembly (<u>assembly FTP</u>) with the HiFi CCS data (hereafter termed “HiFi assembly”) and compared it to a previous FALCON assembly of CHM13 (accession GCA_002884485.1) generated with 77-fold CLR data (hereafter termed “CLR assembly”) (**Fig. 1**). The HiFi assembly required only 2,800 CPU hours, whereas the CLR assembly required more than 50,000 CPU hours. This reduction in runtime is because the N × N correction step common to both FALCON and Canu can be skipped with adequate input read quality. It might be expected that the shorter read length of the HiFi data (N50 10.9 vs. 17.5 kbp) might lead to a less continuous assembly; however, we observed that the HiFi assembly had an N50 of 25.5 Mbp, which is comparable to the N50 of the CLR assembly (29.3 Mbp; **Table 1, Fig. 1**).

**Figure 1.**
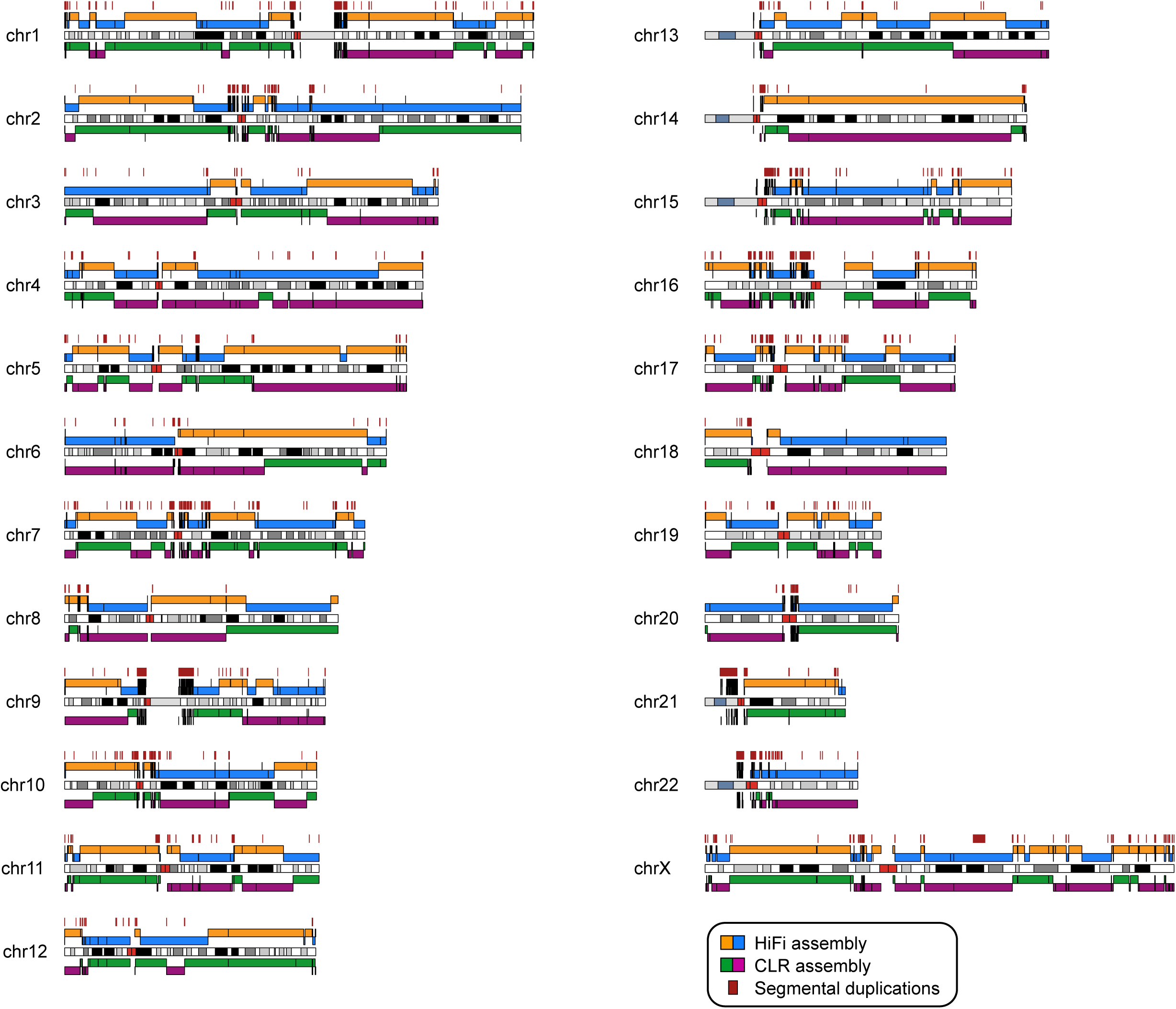
Comparison between the CHM13 HiFi and CLR genome assemblies. Shown are alignments of the HiFi assembly (blue and orange) and the CLR assembly (green and purple) to GRCh38, as well as segmental duplication (SD) blocks greater than 25 kbp in length (dark red) projected onto a karyotype (chromosome banding is indicated in white, black, and gray, with centromeres in bright red and acrocentric regions in blue-gray; CHM13 has a 46X,X karyotype). The alignments are colored by contig name such that when the contig name changes, so does the alignment color. Black bars within a solid color block represent a break in the alignment within the same contig name, which are likely to be locations of structural variants between CHM13 and GRCh38. The large majority of contig alignments over 100 kbp in length end within 50 kbp of an SD [158/166 (95%) in HiFi and 177/182 (97%) in CLR].

To determine assembly base-pair accuracy, we sequenced and assembled the inserts of 31 randomly selected BACs from a genomic library produced from the CHM13 cell line (VMRC59; **Methods**). We estimated assembly accuracy by aligning these sequence inserts to the HiFi and CLR assemblies. We found that, before any polishing, the consensus accuracy of the HiFi assembly was much higher than the CLR assembly (median QV 40.4 vs. 27.5; **Table 1**, **Fig. S2**). Next, we polished the CLR assembly using 77-fold coverage of CLR reads with Quiver and the HiFi assembly using 355-fold coverage of CCS subreads with Arrow. In this experiment, once again, the HiFi assembly was superior to the CLR assembly with respect to accuracy (median QV 43.3 vs. 40.7; **Table 1**, **Fig. S2**).

**Table 1.** Statistics of the HiFi and CLR genome assemblies.

|  | Polishing | Total size (Gbp) | N50 (Mbp) | No. of contigs | Median QV | No. of CPU hours for assembly |
| --- | --- | --- | --- | --- | --- | --- |
| HiFi |  |  |  |  |  |  |
|  | None | 3.03 | 25.51 | 5,296 | 40.41 | ~2,800 |
|  | Arrow | 3.03 | 25.51 | 5,296 | 43.29 | ~10,000 |
|  | Racon | 3.03 | 25.51 | 5,296 | 44.95 | ~2,950 |
|  | 2x Racon | 3.03 | 25.51 | 5,296 | 45.25 | ~3,100 |
|  | 2x Racon+ | 3.03 | 25.51 | 5,296 | 45.25 | ~4,200 |
| CLR |  |  |  |  |  |  |
|  | None | 2.88 | 29.26 | 1,916 | 27.49 | >50,000 |
|  | Quiver | 2.88 | 29.26 | 1,916 | 40.73 | >55,000 |
|  | Quiver+ | 2.88 | 29.26 | 1,916 | 42.70 | >55,000 |

HiFi: HiFi assembly [24-fold sequencing depth] CLR: CLR assembly [72-fold sequencing depth] 2x Racon: Two rounds of Racon 2x Racon+: Two rounds of Racon and one round of Pilon Quiver+: Quiver, Pilon, and FreeBayes-based indel correction Median QV: Median QV over 31 BACs

While the initial assembly of the HiFi data was relatively rapid (2,800 CPU hours), subsequent polishing with Arrow required an additional 7,200 CPU hours. We were curious if we could reduce the polishing time by not incorporating subread information and using only the HiFi data. To do this, we applied Racon (Vaser et al. 2017) to polish our assembly with only the HiFi CCS reads. This Racon-based polishing step finished in only 135 CPU hours (100 for alignment and 35 for polishing) and offered improved accuracy over Arrow (median QV 45.0 vs. 43.3; **Table 1**, **Fig. S2**). After a second round of Racon polishing, there was only one single-nucleotide difference between the HiFi assembly and the BACs excluding indels. Using Illumina WGS data as a third orthogonal platform, we determined that this difference is likely not a sequence error but rather a *bona fide* mutational change that represents a divergence between the propagated VMRC59 BAC and the CHM13 cell line (**Fig. S3**). With the exception of remaining single-base-pair indels, this finding suggests that the QVs reported here should be considered lower bounds due to subsequent propagation errors in BAC DNA (**Supplemental Note**).

To evaluate the global contiguity of the respective assemblies, we generated and applied 2.8-fold sequencing data from strand-specific sequencing (Strand-seq) of the CHM13 cell line. Strand-seq is able to preserve structural contiguity of individual homologs by tracking the read directionality and, therefore, can be used for detection of misassembled contigs in *de novo* assemblies (Falconer et al. 2012; Sanders et al. 2017). Using this analysis, we detected six misassembled contigs that contain seven breakpoints in the HiFi assembly (**Table S1, Fig. S4**). In contrast, we detected a slightly lower number of misassembled contigs (5) and breakpoints (5) in the CLR assembly (**Table S1**). However, given the number of assembled contigs, these results demonstrate that both assemblies are highly accurate, with <0.5% misassembly.

### Segmental duplication analyses

SDs are often recalcitrant to genome assembly due to their high (>90%) sequence identity, length (>1 kbp), and complex modular organization. Therefore, the accuracy and completeness of SDs is a particularly useful metric for assembly quality since these most often correspond to the last gaps in the euchromatic portions of long-read assemblies (Chaisson et al. 2015). We performed a number of analyses to assess the SD resolution in the HiFi and CLR assemblies (**Fig. 2**). First, we compared the percentage of SDs resolved in both genome assemblies, as well as the human reference genome and several recently published long-read assemblies (**Methods**; Vollger et al. 2019). Requiring that SDs are anchored contiguously with unique flanking sequence, we found that, on average, 42% of SDs are resolved in the CHM13 HiFi assembly compared to 32% in the CLR assembly (**Fig. 2A**). Although the majority of human SDs remain unassembled, this is the highest fraction of resolved SDs for any of the published assemblies analyzed thus far (Huddleston et al. 2017; Jain et al. 2018; Seo et al. 2016; Shi et al. 2016), with an average 12% increase over even the ultra-long ONT assembly of NA12878 (**Fig. 2A**). Additionally, the number of bases with significantly elevated coverage (mean + three standard deviations) (Vollger et al. 2019) in the HiFi assembly was reduced by 15% as compared to the CLR assembly (27.3 vs. 32.1 Mbp). This indicates that the HiFi assembly has fewer collapsed sequences compared to the CLR assembly, with multiple SDs now represented by a single contig.

**Figure 2.**
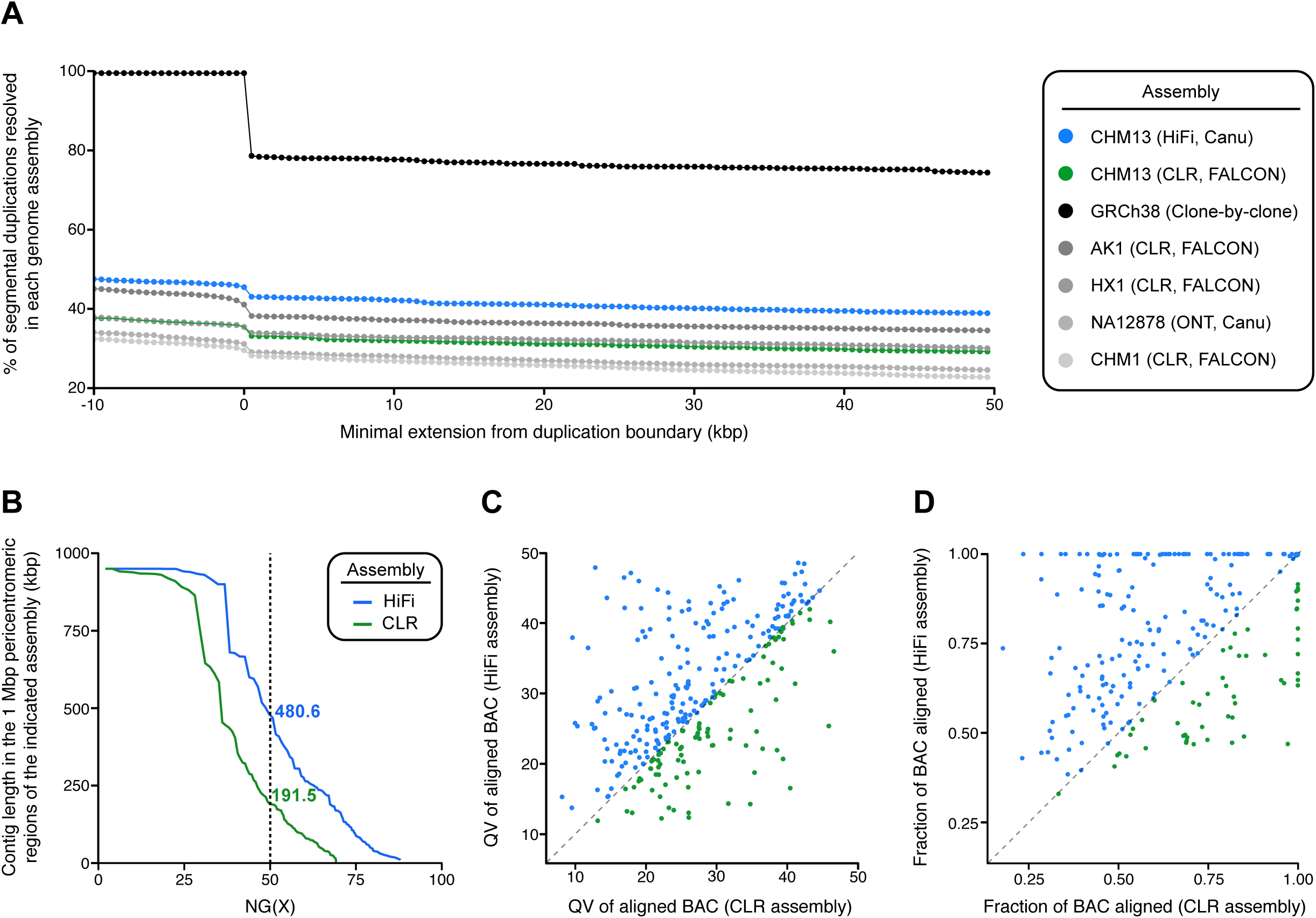
Segmental duplication resolution in the HiFi and CLR genome assemblies. **A)** Shown is the percent of resolved SDs as defined in GRCh38 across the indicated *de novo* assemblies. To be considered resolved, the alignment of the *de novo* assembly must extend X number of base pairs beyond the annotated duplication block on either side. GRCh38 is not 100% resolved after a minimum extension of zero base-pairs because many SDs in GRCh38 are flanked by gaps. **B)** Shown is the NG(X) of the HiFi and CLR assemblies in the 1 Mbp regions flanking the centromeres. NG(X) is defined as the sequence length of the shortest contig at X% of the total pericentromeric region length, which is 46 Mbp (1 Mbp for each pericentromere). The HiFi assembly has an NG50 2.5-fold greater than the CLR assembly in these regions. **C)** Plot of the QV score for each of 310 BACs aligning to SDs within the HiFi and CLR assemblies. Data points above the dashed line have a higher QV score, and, therefore, better sequence identity, in the HiFi assembly relative to the CLR assembly. The accuracy of the HiFi assembly within SDs (median QV 33.5) is increased compared to the CLR assembly (median QV 31.3). **D)** Plot of the fraction of each of 310 BACs aligning to the HiFi and CLR assemblies. Data points above the dashed line have a higher alignment length in the HiFi assembly relative to the CLR assembly. In 253 of the 310 (82%) BACs, the alignment length to the HiFi assembly is greater than or equal to the alignment length in the CLR assembly.

Next, we specifically focused on the pericentromeric regions of the genome where megabases of interchromosomal duplications have accumulated during the course of great ape evolution (She et al. 2004, 2006). We first assessed the contiguity and coverage within the 1 Mbp regions flanking each centromere by calculating a pericentromere-specific NG50. We found that the HiFi assembly had an NG50 of 480.6 kbp, whereas the CLR assembly had a NG50 of only 191.5 kbp (**Fig. 2B**). Next, we assessed contiguity within the pericentromeric regions by counting the number of contigs within the 1 Mbp region flanking the centromeres for each assembly (**Fig. S5A**). Assemblies with fewer contigs have increased contiguity and improved assembly; therefore, we expected that the HiFi assembly would have fewer contigs within many of these regions. Indeed, we found that the HiFi assembly had reduced or the same number of contigs at 52.2% (24/46) of the 1 Mbp pericentromeric regions when compared to the CLR assembly [30.4% (14/46) of the pericentromeric regions had fewer contigs, and 21.7% (10/46) had the same number of contigs in both assemblies]. The remaining pericentromeric regions were split between having no contig representation (8.7%; 4/46) and an increased number of contigs (39.1%; 18/46) in the HiFi assembly relative to the CLR assembly. We hypothesized that the increased number of contigs in these regions in the HiFi assembly may be indicative of fragmented sequences not found in the CLR assembly (**Fig. S5B**). When we tested this hypothesis by summing up the total contig coverage in the 1 Mbp windows flanking the centromeres, we found that, indeed, the HiFi assembly had recovered an additional 5.03 Mbp of pericentromeric sequence missing from the CLR assembly (**Fig. S5C**).

To assess the sequence accuracy and contiguity within SD regions, we compared HiFi and CLR assemblies to 310 sequenced and assembled large-insert BAC clones of CHM13 origin. Once again, we found that the HiFi assembly is more accurate (median QV 33.5, n = 139) than the CLR assembly (median QV 31.3, n = 102) against BACs that align along at least 95% of their length (**Fig. 2C**). We suspect the increased QV is due to the inability of the correction step in FALCON to correctly resolve paralog-specific reads into different groups. Although the HiFi assembly has a higher QV, it should be noted that both assemblies are far less accurate for SDs than unique regions of the genome. Additionally, we find that the HiFi-assembled contigs are more continuous within the sampled SD regions: in 253 of the 310 (82%) BACs, the alignment length to the HiFi assembly is greater than or equal to the alignment length to the CLR assembly (**Fig. 2D**).

A significant fraction of high-identity duplications remain collapsed and unassembled in both the CLR and HiFi assemblies. However, we recently developed a method, Segmental Duplication Assembler (SDA), that can resolve collapsed duplications by taking advantage of long reads that share multiple paralog-specific variants (PSVs) and then grouping them using correlation clustering (Vollger et al. 2019). The algorithm depends on the length of the underlying reads, and since HiFi reads are substantially shorter (N50 10.9 vs. 17.5 kbp), we were concerned that SDA would be limited. To test the ability of HiFi and CLR to resolve collapses, we selected five problematic gene-rich regions of biomedical and biological importance and directly compared the potential of correlation clustering to partition and assemble such regions (**Table S2**; these regions contained the genes *OPN1LW*, *NOTCH2NL*, *SRGAP2*, *FCGR2/3*, *KANSL1*). Of the five regions: two were resolved more accurately by the CLR reads (*OPN1LW, KANSL1*), one was equivalent between HiFi and CLR (*SRGAP2*), and two were better resolved by the HiFi reads (*NOTCH2NL*, *FCGR2/3*). These results are encouraging since SDA was optimized to handle CLR data (Vollger et al. 2019), and we believe future improvements to SDA that take advantage of the high-quality single-nucleotide variants embedded within the HiFi data will resolve even more collapsed regions of genomes.

### Tandem repeat resolution

Since tandem repeat sequences are often difficult to resolve for both length and content, we assessed whether short tandem repeats (STRs) and variable number of tandem repeats (VNTRs) were correctly assembled in the HiFi and CLR assemblies (**Fig. 3**). We identified 3,074 tandem repeats that were ≥1 kbp, on average, across the six Human Genome Structural Variation Consortium (HGSVC) haplotype-resolved assemblies (Chaisson et al. 2019). For each locus, we compared the length of the region in the HiFi and CLR assemblies against an orthogonal set of ultra-long ONT reads generated from CHM13 (**Methods**). A total of 2,969 (96.6%) and 2,936 (95.5%) of the tandem repeats assembled with HiFi and CLR reads, respectively. Both HiFi and CLR assemblies had a high length concordance with ONT reads (Pearson’s correlation coefficients ρ = 0.816 and ρ = 0.809, respectively) over tandem repeats that were resolved in at most a single contig by each assembly and spanned by more than one ONT read (n = 2,898). When we compared loci within each assembly to the mean length of the region in ultra-long ONT reads (with at least one spanning read) (**Fig. 3A**), we found that the HiFi contigs had a lower root-mean-square (RMS) error of 0.886 kbp, while the CLR contigs had an RMS error of 0.952 kbp.

**Figure 3.**
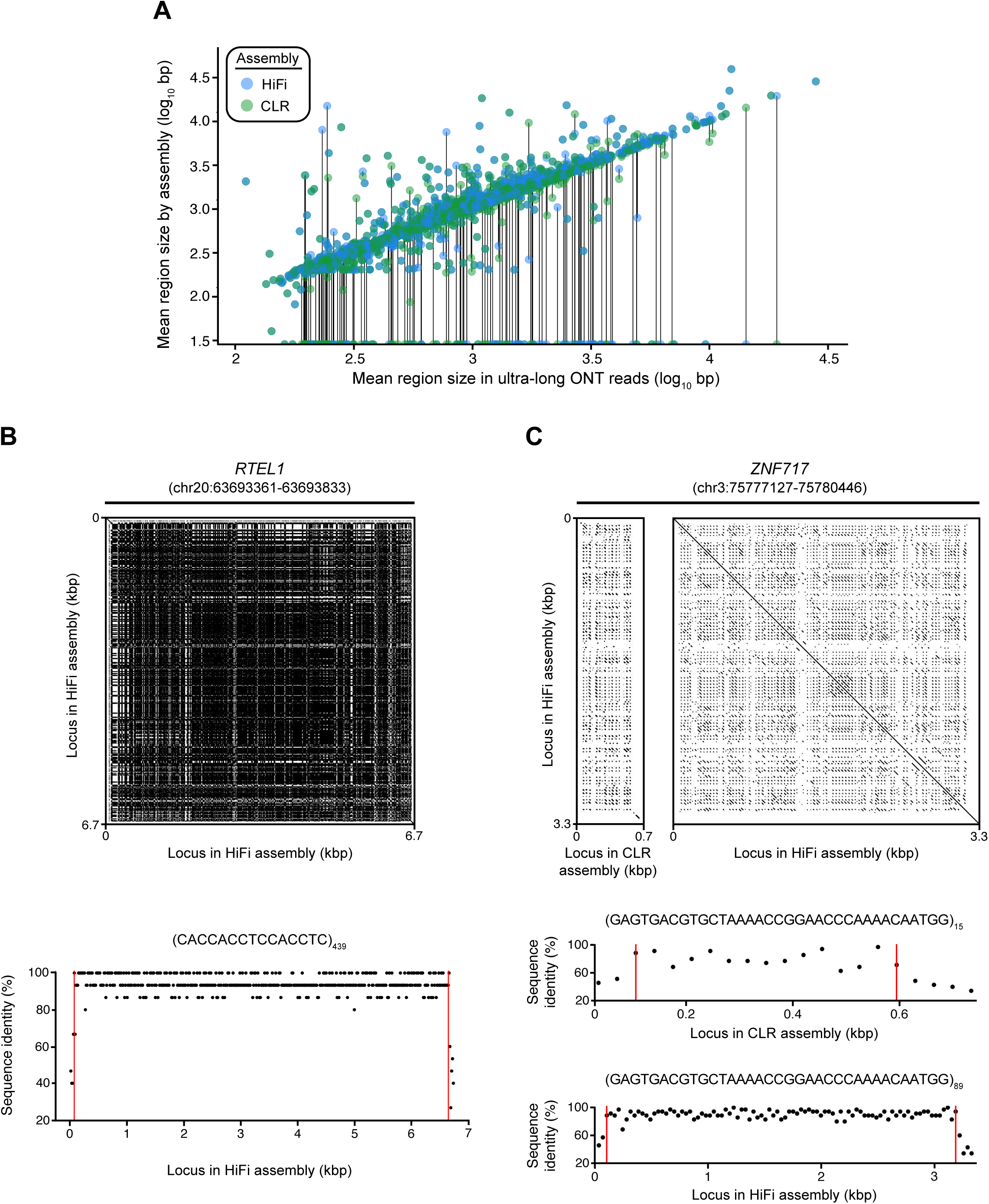
Tandem repeat resolution in the HiFi and CLR genome assemblies. **A)** Plot of the length of tandem repeat loci in the HiFi and CLR assemblies vs. the mean size of these loci in ultra-long CHM13 ONT reads. Discordancy between HiFi and CLR assemblies map off the diagonal, with dropouts clustering as points along on the horizontal axis. For this plot, we include only regions with more than one spanning ONT read and no more than one spanning contig in either assembly (n = 2,898 regions). **B)** Dot plot of a 6.7 kbp VNTR in the intron of *RTEL1* (chr20:63693361-63693833) (top panel), which was resolved in the HiFi assembly only. The CLR assembly contained a gap over this region. The overall structure and length of this VNTR was supported by the ONT reads mapping to this location, which averaged 5,956 +/-1799 bp (n = 5 ultra-long ONT reads), placing the HiFi sequence length at <1 standard deviation away from the average ONT read. The motif homology plot (bottom panel) indicates that the content of the *RTEL1* VNTR is relatively pure, with an average sequence identity to the 15-mer repeat unit of 94.49% across the 439 copies. **C)** Dot plot of the zinc finger protein gene *ZNF717* (GRCh38 coordinates: chr3:75777127-75780446) (top panel), which was collapsed in the CLR assembly but fully represented in the HiFi assembly. The number of copies of this 35 bp repeat unit increased from 15 in the CLR assembly to 89 in the HiFi assembly. The large amount of variation between individual copies of this VNTR is shown in the region between the red lines in the motif homology plots (bottom panels). The level of purity within the VNTR increased from 80.38% sequence identity in the CLR assembly to 90.75% sequence identity in the HiFi assembly. The red vertical lines indicate the start and end position of the VNTR.

Further restricting the analysis to VNTRs present in HiFi but completely absent from the CLR assembly (n = 87), 53% (n = 46) of the loci agreed in length with the ONT reads. Inversely, restricting the analysis to VNTRs present in CLR but completely absent from the HiFi assembly (n = 54), 59% (n = 32) of the loci agreed in length with the ONT reads. The N50 of the 46 validated HiFi-only tandem repeats was 4,968 bp, while the N50 for the 32 validated CLR-only tandem repeats was 3,306 bp. Additionally, the largest VNTRs resolved by HiFi and CLR assemblies were 19,397 bp and 14,250 bp, respectively. This pattern suggests that HiFi reads accurately assemble large tandem repeats that may be inaccessible to CLR. Several of these loci were genic, such as the 439 copy 15-mer in the intron of *RTEL1* (**Fig. 3B**) and the expansion of a 35-mer in the intron of *ZNF717* from 15 (CLR) to 89 (HiFi) tandem repeat copies (**Fig. 3C**). Overall, the HiFi assembly more accurately represented the content and sequence length of the tandem repeats, particularly in previously unrepresented or collapsed regions of the CLR assembly, based on orthogonal validation experiments.

#### Structural variant analyses

Since errors in an assembly will lead to false-positive variant calls, we assessed the utility of assembled HiFi data as a variant discovery tool and used it as a metric to evaluate assembly quality. For each assembly, we called insertions and deletions against GRCh38 from contig alignments and filtered for consensus regions (loci where the assembly had one mapped contig, **Methods**). We generated a callset for each assembly before and after polishing using a variety of tools, including Racon, Quiver, Arrow, Pilon, and a FreeBayes-based indel correction pipeline (Chin et al. 2013; Kronenberg et al. 2018; Vaser et al. 2017; Walker et al. 2014). We found that SV (indels ≥50 bp) calls were largely consistent among assemblies (**Table 2**). Although HiFi read quality is substantially higher, polishing was required to reduce the number of false positive indel calls (**Table 3**). Overall, we found that the number of insertions and deletions were comparable between polished HiFi and CLR assemblies. When we compare SVs to published CHM13 calls, we see very strong concordance, with 89.5% of insertions and 86.8% of deletions called in both (**Fig. S6**).

**Table 2.** Summary of SV calls in the HiFi and CLR assemblies.

| Polishing | Insertions |  |  | Deletions |  |  | All |  |  |
| --- | --- | --- | --- | --- | --- | --- | --- | --- | --- |
|  | No. of events | Mean length (bp) | Total length (bp) | No. of events | Mean length (bp) | Total length (bp) | No. of events | Mean length (bp) | Total length (bp) |
| HiFi |  |  |  |  |  |  |  |  |  |
| None | 10,650 | 569 | 6,063,301 | 6,254 | 482 | 3,012,598 | 16,904 | 537 | 9,075,899 |
| Arrow | 10,608 | 570 | 6,050,398 | 6,243 | 482 | 3,009,027 | 16,851 | 538 | 9,059,425 |
| Racon | 10,632 | 569 | 6,044,348 | 6,273 | 478 | 3,000,394 | 16,905 | 535 | 9,044,742 |
| 2x Racon | 10,603 | 570 | 6,048,876 | 6,273 | 479 | 3,005,723 | 16,876 | 537 | 9,054,599 |
| 2x Racon+ | 10,579 | 571 | 6,044,564 | 6,468 | 475 | 3,072,543 | 17,047 | 535 | 9,117,107 |
| CLR |  |  |  |  |  |  |  |  |  |
| None | 10,655 | 558 | 5,947,788 | 6,405 | 471 | 3,018,503 | 17,060 | 526 | 8,966,291 |
| Quiver | 10,664 | 559 | 5,959,522 | 6,497 | 476 | 3,095,325 | 17,161 | 528 | 9,054,847 |
| Quiver+ | 10,627 | 560 | 5,950,702 | 6,992 | 469 | 3,275,883 | 17,619 | 524 | 9,226,585 |

HiFi: HiFi assembly CLR: CLR assembly 2x Racon: Two rounds of Racon 2x Racon+: Two rounds of Racon and one round of Pilon Quiver+: Quiver, Pilon, and FreeBayes-based indel correction SV: indel ≥ 50 bp Excludes SVs mapping to pericentromeric regions (see **Methods**)

**Table 3.** Summary of indels in the HiFi and CLR assemblies.

| Polishing | Insertions |  |  |  | Deletions |  |  |  |
| --- | --- | --- | --- | --- | --- | --- | --- | --- |
|  | No. of events | % of 1 bp events | Mean length (bp) | Total length (bp) | No. of events | % of 1 bp events | Mean length (bp) | Total length (bp) |
| HiFi |  |  |  |  |  |  |  |  |
| None | 1,014,192 | 80% | 1.87 | 1,891,600 | 1,340,486 | 86% | 1.69 | 2,269,778 |
| Arrow | 339,429 | 50% | 3.46 | 1,172,772 | 345,678 | 48% | 3.66 | 1,264,044 |
| Racon | 343,927 | 50% | 3.44 | 1,182,117 | 342,106 | 48% | 3.67 | 1,254,412 |
| 2x Racon | 343,733 | 50% | 3.44 | 1,181,197 | 339,831 | 48% | 3.68 | 1,251,757 |
| 2x Racon+ | 343,132 | 50% | 3.44 | 1,179,652 | 339,787 | 48% | 3.69 | 1,252,463 |
| CLR |  |  |  |  |  |  |  |  |
| None | 943,936 | 75% | 1.99 | 1,860,855 | 3,616,964 | 82% | 1.46 | 5,271,942 |
| Quiver | 350,924 | 50% | 3.41 | 1,196,221 | 509,229 | 62% | 2.87 | 1,460,402 |
| Quiver+ | 353,245 | 50% | 3.39 | 1,198,153 | 392,657 | 52% | 3.41 | 1,337,638 |

HiFi: HiFi assembly CLR: CLR assembly 2x Racon: Two rounds of Racon 2x Racon+: Two rounds of Racon and one round of Pilon Quiver+: Quiver, Pilon, and FreeBayes-based indel correction

### Gene open reading frame annotations

Long-read sequencing platforms exhibit high indel error rates due to missed and erroneous incorporations during real-time sequencing. As a result, predicted open reading frames are often disrupted, leading to potential problems in gene annotation (Watson and Warr 2019) unless additional error correction steps are employed (Kronenberg et al. 2018). We compared the SV and indel callsets to human RefSeq annotations and identified likely gene-disruptive events (**Methods**). In the unpolished HiFi assembly, we found 16,158 SVs and indels putatively disrupting 4,151 of 18,045 RefSeq genes within the assembly consensus regions (23%), which reduced to 134 after polishing with two rounds of Racon (0.74%) (**Table 4**). Before polishing, these predicted gene-disruptive SVs and indels were overwhelmingly single-base-pair errors (98%; 15,822 of 16,158), which were greatly reduced after polishing (56%; 93 of 165). As expected, the CLR assembly had more likely-disrupted genes before polishing (64%; 11,593 of 17,991 genes in its consensus region), but this declined to 209 after polishing (1.2%). We found fewer predicted disrupted genes outside of repetitive events in the HiFi assembly (53 in HiFi vs. 58 in CLR), and this trend increases inside SDs where short reads may not polish as effectively (39 in HiFi vs. 101 in CLR). It is worth noting that 2,412 protein-coding genes (13%) have exons in SDs, and this difference between the HiFi and CLR assemblies represents 2.7% of these duplicated protein-coding genes.

**Table 4.** Summary of disrupted RefSeq gene models in the HiFi and CLR assemblies.

| Polishing | No. of events in whole genome | No. of events in whole genome excluding TRs/SDs | No. of events in SDs only |
| --- | --- | --- | --- |
| HiFi |  |  |  |
| None | 4,151 | 2,481 | 360 |
| Arrow | 138 | 55 | 39 |
| Racon | 154 | 65 | 40 |
| 2x Racon | 135 | 54 | 39 |
| 2x Racon+ | 134 | 53 | 39 |
| CLR |  |  |  |
| None | 11,593 | 6,686 | 1,249 |
| Quiver | 653 | 261 | 159 |
| Quiver+ | 209 | 58 | 101 |

HiFi: HiFi assembly CLR: CLR assembly 2x Racon: Two rounds of Racon 2x Racon+: Two rounds of Racon and one round of Pilon Quiver+: Quiver, Pilon, and FreeBayes-based indel correction No. of events in whole genome: All RefSeq gene models within the assembly consensus regions were counted. Total gene count is 18,045 (HiFi assembly) and 17,991 (CLR assembly). No. of events in whole genome excluding TRs/SDs: All RefSeq gene models within the assembly consensus were counted except for those with exons intersecting tandem repeats (TRs) or segmental duplications (SDs). Total gene count is 10,853 (HiFi assembly) and 10,850 (CLR assembly). No. of events in SDs only: Only RefSeq gene models within SDs were counted. Total gene count is 2,005 (HiFi assembly) and 1,951 (CLR assembly).

Since true biological variation and reference errors will contribute to gene-disrupted events, we expect many of these to be biological and not necessarily assembly artifacts. When we intersect the disrupted genes from the polished HiFi and CLR assemblies, we find that the HiFi genes are largely a subset of the CLR genes, but the converse is not true (**Fig. S7**). To provide additional support for these events, we intersected gene-disrupting variants with CHM13 calls from SMRT-SV (Audano et al. 2019) and a FreeBayes callset from Illumina CHM13 whole-genome sequence reads (ERR1341795) (**Methods**). We applied this to both the polished HiFi assembly (2 times with Racon) and the fully polished CLR assembly. In the HiFi assembly, 13% (17 of 135) of the disrupted genes had no orthogonal support with the majority corresponding to duplicated genes (14 genes). We conclude that the events in these 17 genes are likely false positives; however, only three of these remaining unsupported gene-disrupting indels mapped to unique sequence. In the CLR assembly, 44% (93 of 209) of the gene-disruptive events had no orthogonal support with the majority (80 genes) mapping to SDs. These experiments suggest that there are approximately 120 genes in CHM13 altered by *bona fide* frame-shifting indels and SVs when compared to GRCh38 and RefSeq annotations.

## Discussion

The generation and assembly of HiFi and CLR long-read sequence data from the same haploid source material allows us to directly compare the accuracy and contiguity of these technologies without the added complication of disentangling haplotypes needed to resolve SV alleles. We conclude that there are three key strengths of the HiFi technology over CLR technology. First, the time to generate the *de novo* assembly is reduced 10-fold, and it will likely be reduced further as HiFi assemblers are developed and optimized. This not only makes *de novo* assembly of human genomes accessible to a larger number of research groups, but it also paves the ways for larger cohorts of individuals to be sequenced and assembled. Although assembly time is drastically reduced, the background compute time required to generate HiFi data by the CCS algorithm remains substantial (∼50,000 CPU hours in total).

Second, our analyses confirm that, both in terms of quality and continuity, the HiFi assembly is generally superior or at least comparable to the CLR assembly despite the shorter read lengths and effectively reduced genome coverage (Wenger et al. 2019). One significant advance is that HiFi assembly can be polished without reverting to the underlying subreads, which saves approximately 1 terabyte of subread data and 7,000 hours of additional compute time. Polishing remains an absolute requirement to reduce indel errors and obtain a high-quality final assembly. Human CLR datasets ultimately require orthogonal Illumina data, and our results show that the HiFi sequencing platform alone achieves a greater level of accuracy for annotated protein-coding genes.

Finally, we demonstrate that, in some of the most difficult regions of the genome (i.e., SDs, pericentromeric regions, and tandem repeats), the HiFi assembly shows improved continuity and representation, but relatively modest accuracy improvements (**Figs. 1-3, Fig. S5, and Tables 1-4**). Highly accurate HiFi data allows for the assembly of an additional 10% of duplicated sequences and better recovers the structure of tandem repeats such that they more exactly reflect the genomic length of VNTRs and STRs as confirmed by orthogonal analyses. We note, however, that the accuracy of the duplicated and tandem repeat regions is still lower than that of unique regions of the genome. Follow-up procedures such as SDA, which are designed to target and further resolve collapsed regions, show mixed results especially among the most highly identical human duplications. Our analyses suggest that this is a limitation of the shorter read lengths of HiFi (N50 of 10.9 vs. 17.5 kbp), which reduces the power needed to phase PSVs and assign collapsed reads to their respective duplicated loci. Nevertheless, we believe the results are encouraging since methods such as SDA were optimized to handle CLR data (Vollger et al. 2019). Future improvements to SDA that take advantage of the high-quality single-nucleotide variants embedded within the HiFi data in duplicated regions will resolve even more collapsed regions of assembled genomes.

Next steps involve benchmarking and optimization of performance within diploid genome assemblies. Much of the recent advances in improving the contiguity of genome assemblies from telomere to telomere (Miga, Koren et al., unpublished) have been based on the same haploid source material analyzed here. It is clear that current HiFi genome assemblies are not as contiguous as those generated with high-coverage, ultra-long ONT data, or with combinations of PacBio and ONT data. While the haploid source material has been extremely useful for benchmarking, the ultimate challenge is the accurate assembly of human diploid genomes where both chromosomal haplotypes are fully resolved. Incorporation of linking-read technologies, such as Strand-seq, Hi-C, and 10X Genomics, or trio-binning approaches have been shown to significantly improve phasing and SV sequence and assembly (Koren et al. 2018; Chaisson et al. 2019; Kronenberg et al. 2019). It is likely that such approaches could be combined with HiFi datasets to enhance telomere-to-telomere phasing and improve the accuracy of more complex repeats. Alternatively, the use of ultra-long-read datasets coupled with HiFi sequencing on the same samples will likely enhance both the phasing and accuracy of diploid genome assemblies. A useful standard for diploid genome assembly will be to repeat these analyses for two haploid source genomes in order to model the effect and accuracy of *in silico* diploid genomes as we (Huddleston et al. 2017) and others (Li et al. 2018) have shown.

Notwithstanding these advances, significant challenges remain for complete genome assembly, including large SDs, centromeric satellites, and acrocentric regions. For example, although the CHM13 HiFi assembly we generated is highly contiguous (N50 25.5), an analysis of the unmappable reads shows an abundance of repetitive DNA (70.4%; **Fig. S8**). Of these sequences, 49.5% consist of various classes of satellite repeats, which populate centromeres and the acrocentric portions of human chromosomes. Given the accuracy of these unmapped sequence reads, they will be quite valuable in obtaining the first overview of the sequence content and composition of these more complex heterochromatic regions. Obtaining even longer HiFi reads than used in this assembly (i.e., >11 kbp average used here) will be necessary to accurately anchor and sequence-resolve these repeat regions in future genome assemblies. Coupled with advances from other long-read technologies, such as ONT, it is clear that highly accurate telomere-to-telomere assemblies of diploid genomes will soon be achievable.

## Methods

### Cell lines

Cells from a complete human hydatidiform mole, CHM13 (46X,X), were immortalized with human telomerase reverse transcriptase (hTERT) and cultured in complete AmnioMAX C-100 Basal Medium (ThermoFisher Scientific, Carlsbad, CA) supplemented with 15% AmnioMAX supplement (ThermoFisher Scientific, Carlsbad, CA) and 1% penicillin and streptomycin. Cells were maintained at 37°C in a humidified incubator with 5% CO_2_.

### CCS library preparation

High-molecular-weight DNA was isolated from cultured CHM13 cells using a modified Qiagen Gentra Puregene Cell Kit protocol (Huddleston et al. 2014). A HiFi library with an average insert length of ∼11 kbp was generated according to the protocol in Wenger et al. 2019 and sequenced on four SMRT Cells 8M on a Sequel II instrument using Sequel II Sequencing Chemistry 1.0, 12-hour pre-extension, and 30-hour movies. Raw data was processed using the CCS algorithm (version 3.4.1, parameters: --minPasses 3 --minPredictedAccuracy 0.99 --maxLength 21000) to yield 75.7 Gbp in 6.9 million reads with an average read length of 10.95 kbp and estimated median QV of 32.85. Sequence data is available via NCBI SRA (https://www.ncbi.nlm.nih.gov/sra/SRX5633451). Average run time for the CCS algorithm was ∼12,500 CPU core hours per SMRT Cell (∼50,000 total).

### Strand-seq library preparation

Cultured CHM13 cells were pulsed with BrdU and used for preparation of single-cell Strand-seq libraries as previously described (Sanders et al. 2017).

### BAC clone insert sequencing

BAC clones from the VMRC59 clone library were hybridized with probes targeting complex or highly duplicated regions of GRCh38 (n = 310), or selected from random regions of the genome not intersecting with SD (n = 31). DNA from positive clones was isolated, screened for genome location, and prepared for long-insert PacBio sequencing as previously described (Vollger et al. 2019). Libraries were sequenced on the PacBio RS II and Sequel platforms with the P6-C4 or Sequel 2.1/Sequel 3.0 chemistries, respectively. We performed *de novo* assembly of pooled BAC inserts using Canu v1.5 (Koren et al. 2017). After assembly, we removed vector sequence (pCCBAC1), restitched the insert, and then polished with Quiver or Arrow. Canu is specifically designed for assembly with long error-prone reads, whereas Quiver/Arrow is a multi-read consensus algorithm that uses the raw pulse and base call information generated during SMRT sequencing for error correction. We reviewed PacBio assemblies for misassembly by visualizing the read depth of PacBio reads in Parasight (http://eichlerlab.gs.washington.edu/jeff/parasight/index.html), using coverage summaries generated during the resequencing protocol.

### Genome assembly

Canu v1.7.1 was applied with the following parameters to generate the HiFi *de novo* assembly: `genomeSize=3.1g correctedErrorRate=0.015 ovlMerThreshold=75 batOptions=“-eg 0.01 -eM 0.01 -dg 6 -db 6 -dr 1 -ca 50 -cp 5” -pacbio-corrected`

Assemblies were mapped to GRCh38 with minimap2 (Li 2017) version 2.15 using the following parameters: `--secondary=no -a --eqx -Y -x asm20 -m 10000 -z 10000,50 -r 50000 --end-bonus=100 -O 5,56 -E 4,1 -B 5`. These alignments were used for downstream SV calling and ideogram visualizations.

Error correction with Quiver, Arrow, Pilon, and indel correction was done as previously described (Chin et al. 2013; Kronenberg et al. 2018; Vaser et al. 2017; Walker et al. 2014). Error correction with Racon was executed with the following steps:

`minimap2 -ax map-pb --eqx -m 5000 -t {threads} --secondary=no {ref} {fastq} | samtools view -F 1796 - > {sam}`

`racon {fastq} {sam} {ref} -u -t {threads} > {output.fasta}`

### QV calculations

QV calculations were made by alignments to 31 sequenced and assembled BACs falling within unique regions of the genome (>10 kbp away from the closest SD) where at least 95% of the BAC sequence was aligned. The following formula was used to calculate the QV, and gaps of size N were counted as N errors: QV = –10log_10_[1 – (percent identity/100)]. QV calculations within SDs were done in the same manner but against 310 BACs that overlap with SD regions.

### SD analyses

SDs were defined as resolved or unresolved based on their alignments to GRCh38 using the minimap2 parameters described above. Alignments that extended a minimum number of base pairs beyond the annotated SDs were considered to be resolved. This minimum extension varied from -10,000 to 50,000 bp and the average difference between assemblies was used to define the percent difference reported.

The number of collapsed bases was determined by aligning the CLR reads to both the CLR and the HiFi assemblies. Regions were defined as collapsed if they met the following conditions: coverage greater than the mean coverage plus three standard deviations, 15 kbp of consecutive increased coverage or more, and <80% repeat content as defined by RepeatMasker.

### Pericentromeric analyses

The number of contigs within each pericentromeric region was calculated by first aligning the contigs from the HiFi or CLR assemblies to GRCh38 using the minimap2 parameters described above. Alignments were limited to be within 1 Mbp on either side of the centromere decoys, and then unique contig names were counted.

The representation within the pericentromeric regions was calculated using bedtools to collapse all filtered contigs within the pericentromeric region for the HiFi and CLR assemblies. The resulting size of the collapsed contigs within the CLR assembly was subtracted from the size calculated in the corresponding region in the HiFi assembly.

The pericentromere-specific NG50 statistic was calculated using a G of 46 Mbp (accounting for the 1 Mbp size of each pericentromeric region on the 23 chromosomes).

### Tandem repeat analyses

Tandem Repeats Finder (Benson 1999) was run on the six haplotype-resolved assemblies (Chaisson et al. 2019) as well as the CLR CHM13 assembly using the following parameters: 2 7 7 80 10 50 2000 -h -d -ngs. After identifying all tandem repeats not represented or collapsed in the CLR assembly relative to the six human haplotypes, we obtained a final set of 3,074 large tandem repeats, all of which were anchored in GRCh38. Second, we retrieved sequence from each of these loci using the two assemblies and our orthogonal CHM13 ONT data source. For each region in both assemblies and aligned ultra-long ONT reads, we extracted the sequence that mapped from the start of the region to the end using the alignment CIGAR strings as a guide. Since multiple sequences may map to a region, we recorded the number of alignments and computed the average length of the region for each dataset. Concordance with ONT reads was defined by allowing ≤5% variation in the average ONT read length. For our in-depth sequence analysis of the two VNTR loci, we used repeat homology plots, which were constructed using a pairwise alignment between the motif and assembled sequence in every tiling window of the same length as the repeat unit length (i.e., 15 bp and 53 bp, respectively, for the two VNTRs; **Figs. 3B,C**). At any given window, the repeat unit (i.e., the motif) was circularized in 1 bp increments, and the maximal sequence identity was reported at each tiling window. The dotplots were generated using Gepard (Krumsiek, Arnold, and Rattei 2007).

### SV analyses

For assembly in each polishing stage, contigs mapped to GRCh38 were used to create a consensus region, which included all loci with exactly one aligned contig. Next, we called indels and SVs from the alignments using a previously validated method (Chaisson et al. 2015) implemented in PrintGaps.py distributed in the SMRT-SV v2 pipeline (https://github.com/EichlerLab/smrtsv2). We then filtered for variants within the assembly’s consensus region. We further filtered out variants in pericentromeric loci where callsets are difficult to reproduce (Audano et al. 2019). This process was repeated for each assembly in each polishing stage.

For gene annotations, SVs were intersected with a callset from SMRT-SV and FreeBayes. For the SMRT-SV indels, we retrieved the CHM13 contigs, called SVs and indels from them using the same PrintGaps.py method. SMRT-SV generates a BED file linking regions of GRCh38 to the best contig for variant calling, and we used this BED to filter the SV and indel calls from the overlapping assembly contigs. We then intersected HiFi and CLR variants with either SMRT-SV or FreeBayes SVs and indels using custom code that requires either a variant length match by 50% and maximum distance between events is no more than 50 bp or 50% reciprocal overlap. Matching by size and distance reduces overlap bias for short indels while matching by reciprocal overlap allows larger SVs to intersect even when they are shifted, which is common for calling insertions associated with tandem duplications or repetitive sequence.

### Gene annotation

With custom code using the SV and indel callset, the number of bases in coding regions of RefSeq annotations (retrieved 2019-04-24 from UCSC RefSeq track on GRCh38) were quantified. Briefly, if an insertion was located in a coding region, its entire length was taken as the number of coding bases it affects. For deletions, the number of bases falling inside the coding region were quantified. From these results, we obtained a set of genes where at least one variant inserts or deletes a number of bases that is not a multiple of three within any isoform of the gene. For this analysis, we excluded RefSeq noncoding RNA annotations.

We intersected RefSeq exons with tandem repeats (UCSC hg38 “Simple Repeats” track) and SDs (UCSC hg38 “Segmental Dups” track) to annotate them as either containing or absent of SDs or tandem repeats. For each assembly, we calculated results using only RefSeq genes that are fully contained within its consensus region.

### RepeatMasker analysis of unmappable sequences

All HiFi sequence reads were mapped to the *de novo* assemblies using the following minimap2 parameters: -x asm20 -m 4000 --secondary=no --paf-no-hit. Reads that did not map to the *de novo* assemblies were subjected to RepeatMasker analysis (Smit, Hubley, and Green 1996) to determine their repeat content.

### Data Access

HiFi assemblies with varying levels of polishing are available here: https://eichlerlab.gs.washington.edu/help/mvollger/papers/chm13_hifi/rebasecalled/HiFi_Asms/. CLR assemblies with varying levels of polishing are available here: https://eichlerlab.gs.washington.edu/help/mvollger/papers/chm13_hifi/rebasecalled/CLR_Asms/. HiFi sequence data (SRX5633451), CLR sequence data (SRX818607, SRX825542, and SRX825575-SRX825579), and assembled BACs from the VMRC59 clone library are available via NCBI SRA.

## Acknowledgments

The authors thank T. Brown for assistance in editing this manuscript. This work was supported, in part, by grants from the U.S. National Institutes of Health (NIH grants HG002385 and HG010169 to E.E.E.) and an Advanced Grant from the European Research Council (P.M.L.). M.R.V. was supported by a National Library of Medicine (NLM) Big Data Training Grant for Genomics and Neuroscience (5T32LM012419-04). A.S. was supported by a National Human Genome Research Institute (NHGRI) Training Grant (5T32HG000035-23). E.E.E. is an investigator of the Howard Hughes Medical Institute.

## Author Contributions

M.R.V., G.A.L., and E.E.E. wrote the manuscript; M.R.V., G.A.L., P.A.A., A.S., and D.P. produced the display items; M.R.V. performed the assembly and polishing with suggestions from Z.N.K. and A.M.W.; M.R.V. and A.M.W. performed the QV analysis; D.P. performed the Strand-seq analysis; M.R.V. and G.A.L. performed the SD analyses; A.S. and P.A.A. performed the tandem repeat analysis; P.A.A. performed the SV and gene annotation analyses; G.A.L. performed the unassembled sequence analysis; M.R.V. and G.A.L. organized the supplementary material; P.P., G.T.C., K.M.M., C.B., and M.W.H. generated the PacBio genome sequence data; U.S. developed and supplied the homozygous CHM13hTERT cell line.

## Supplemental material for Vollger, Logsdon et al

### Supplemental Figure Legends

**Figure S1.**
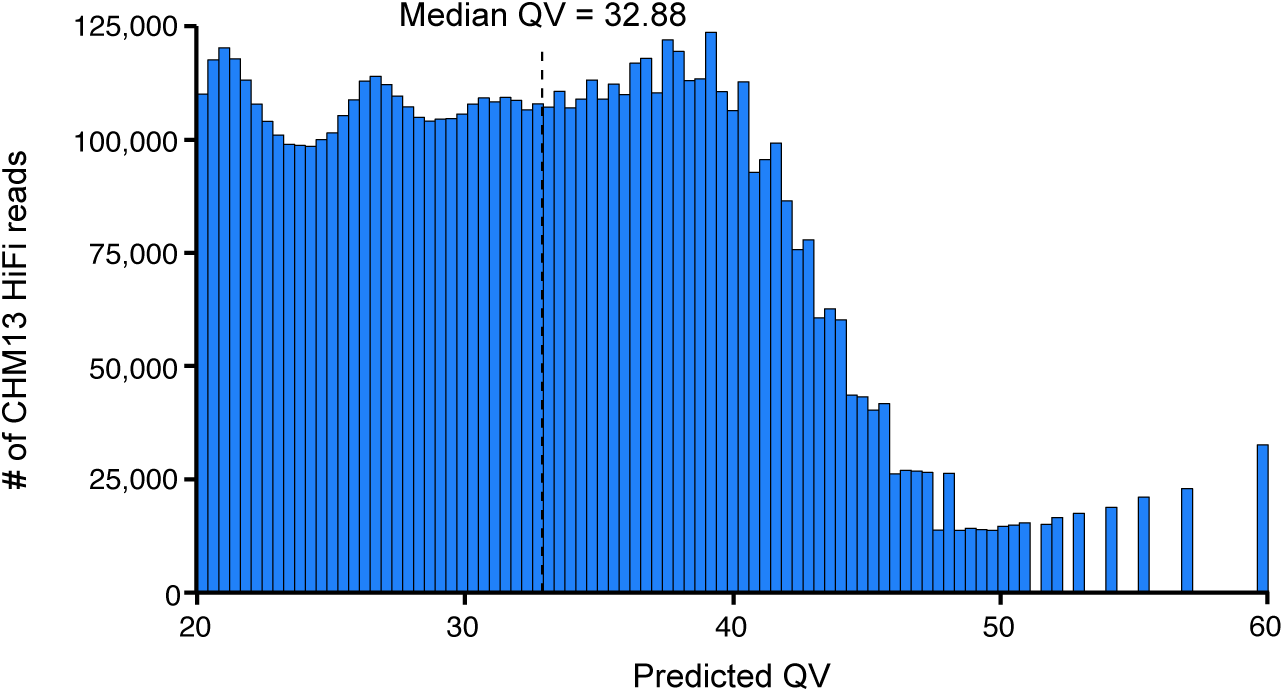
Distribution of read quality value (QV) within the CHM13 HiFi dataset. Plot of the PacBio-predicted read qualities for all HiFi data with a QV > 20. The median QV of this data is 32.88. This dataset was used for *de novo* assembly and all other analyses.

**Figure S2.**
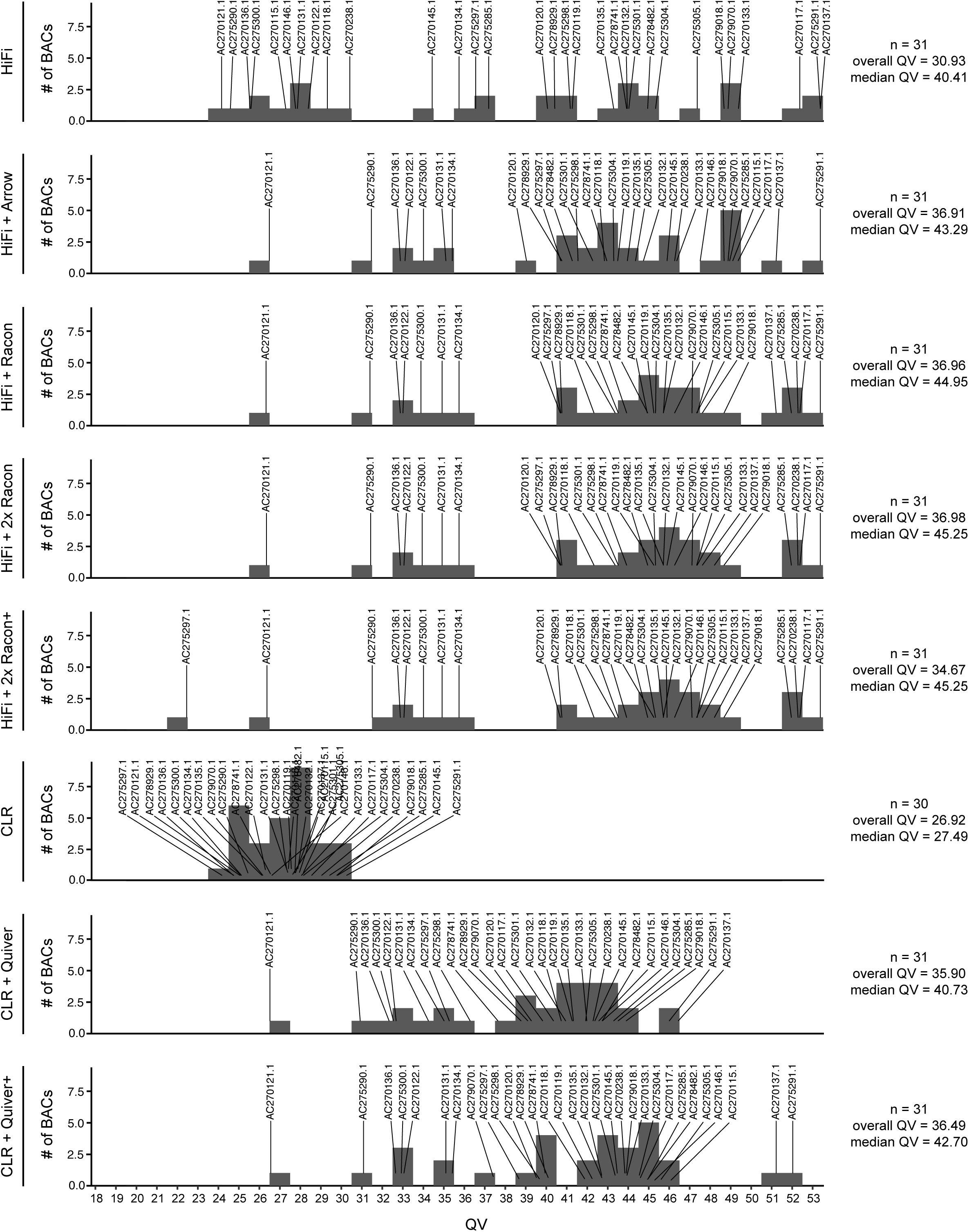
Assessment of QV score of each genome assembly with varying levels of polishing. Histogram of QV score derived from the alignment of 31 BACs to the indicated assembly. Each BAC clone accession name is indicated, and the overall and median QV scores for each genome assembly is shown.

**Figure S3.**
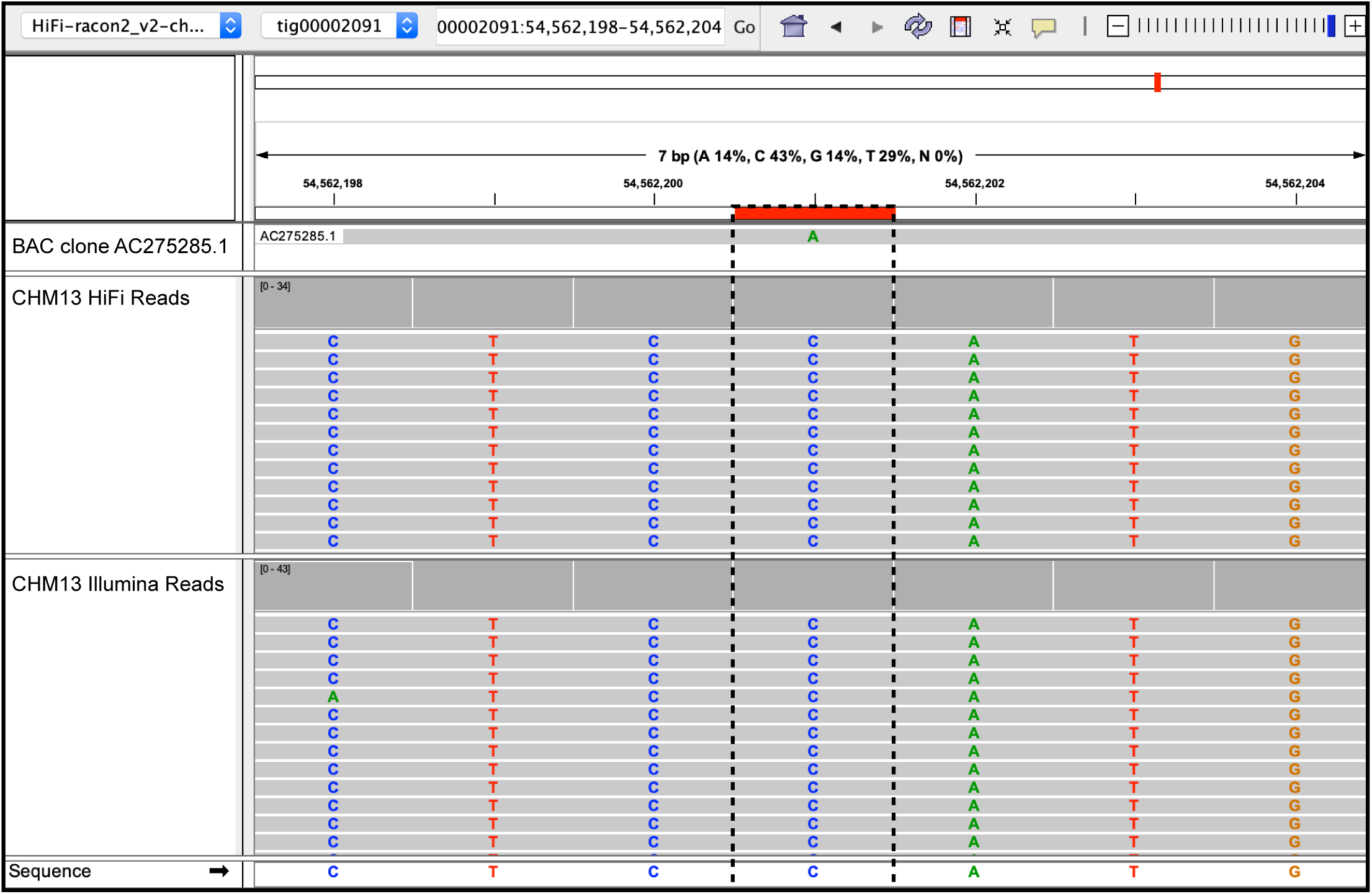
Assessment of a base mismatch between a BAC clone and the HiFi assembly. Out of all 31 BACs assessed, there was a single mismatch observed between one BAC and the HiFi assembly (highlighted here in red and boxed with a dashed line). However, both the HiFi and Illumina data supported the base in the HiFi assembly (a ‘C’ rather than an ‘A’), indicating a divergence in sequence between BAC clone AC275285.1 and the CHM13 cell line (see **Supplemental Notes**).

**Figure S4.**
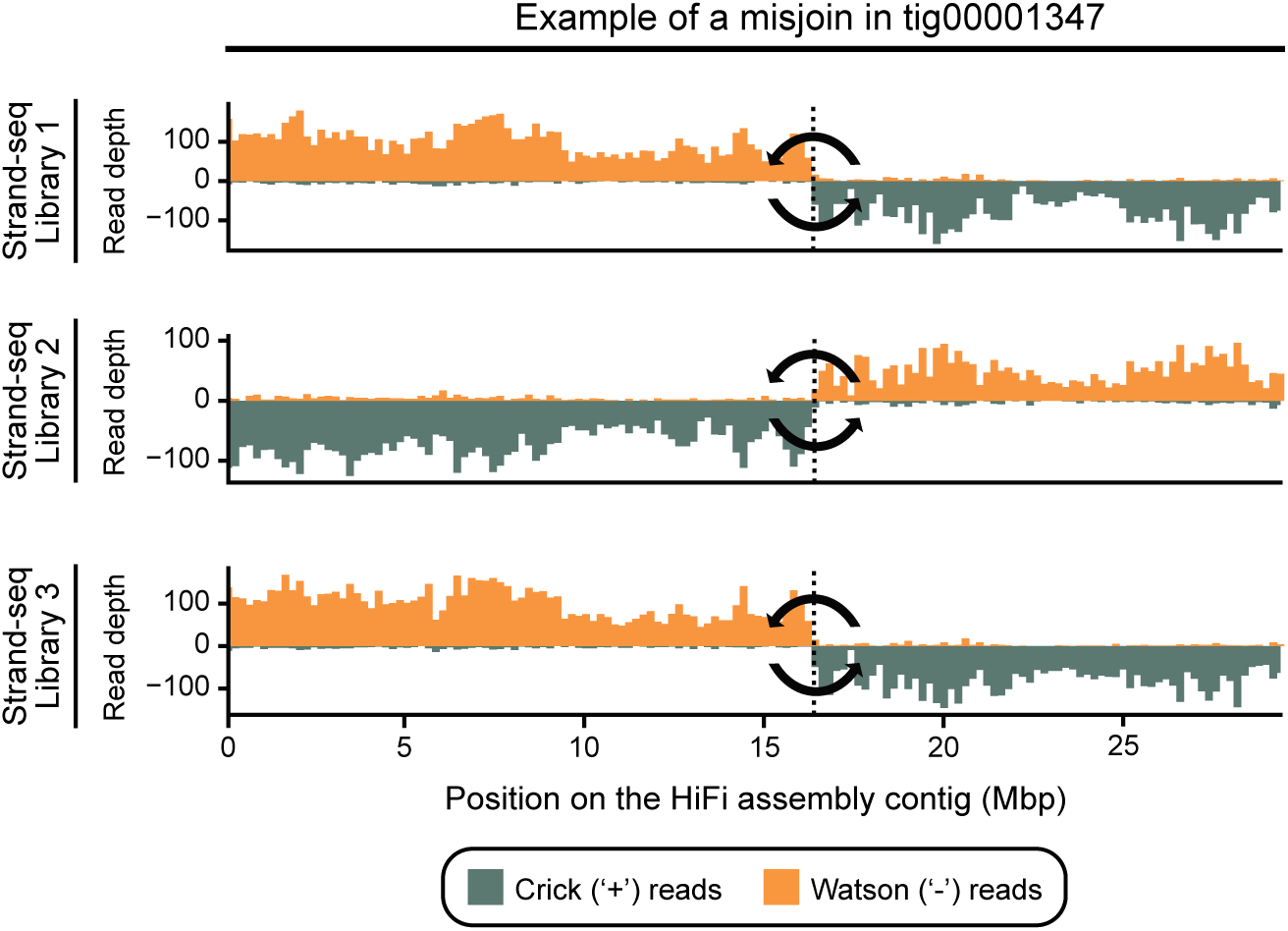
Example of a misjoined contig in the HiFi assembly. Shown is an example of a misjoined contig in the HiFi assembly. Reads mapping to the plus (Crick; teal) or minus (Watson; orange) strand of the reference genome are plotted as vertical bars along the contig. Each row shows one Strand-seq library. A recurrent change in read directionality in the middle of the contig suggests that left and right portions of this contig have flipped orientation with respect to each other and have likely been misjoined during the assembly process.

**Figure S5.**
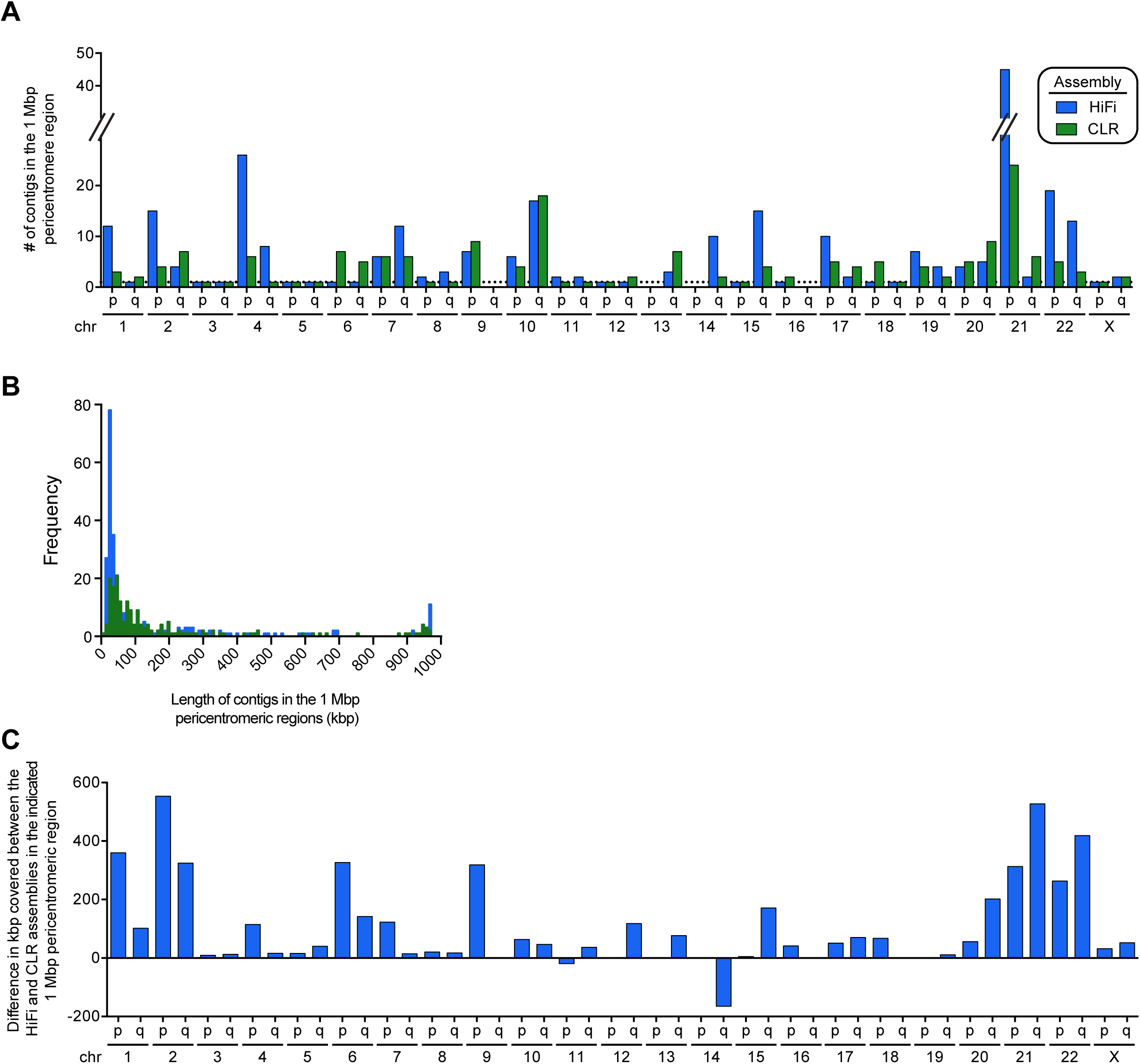
Assessment of continuity in the pericentromeric regions in the HiFi and CLR assemblies. **A)** Plot of the number of contigs in the 1 Mbp regions flanking each centromere in the HiFi and CLR assemblies. The majority of the pericentromeric regions in the HiFi assembly (52.2%) contained either a reduced number of contigs or the same number of contigs. The remaining pericentromeric regions either contained no contig (8.7%) or an increased number of contigs (39.1%) in the HiFi assembly relative to the CLR assembly. **B)** Histogram of the length of contigs in the 1 Mbp regions flanking the centromeres for each assembly. The HiFi assembly has more contigs than the CLR assembly overall, with an increase in the number of small contigs (<100 kbp) and large contigs (900-1000 kbp). The average contig length is 145.8 kbp in the HiFi assembly and 177.6 kbp in the CLR assembly. **C)** Plot of the difference in sequence coverage in the 1 Mbp regions flanking each centromere for the HiFi and CLR genome assemblies. Nearly all pericentromeric regions contain additional sequences in the HiFi assembly relative to the CLR assembly. The HiFi assembly contains an additional 5.03 Mbp of pericentromeric sequence missing in the CLR assembly.

**Figure S6.**
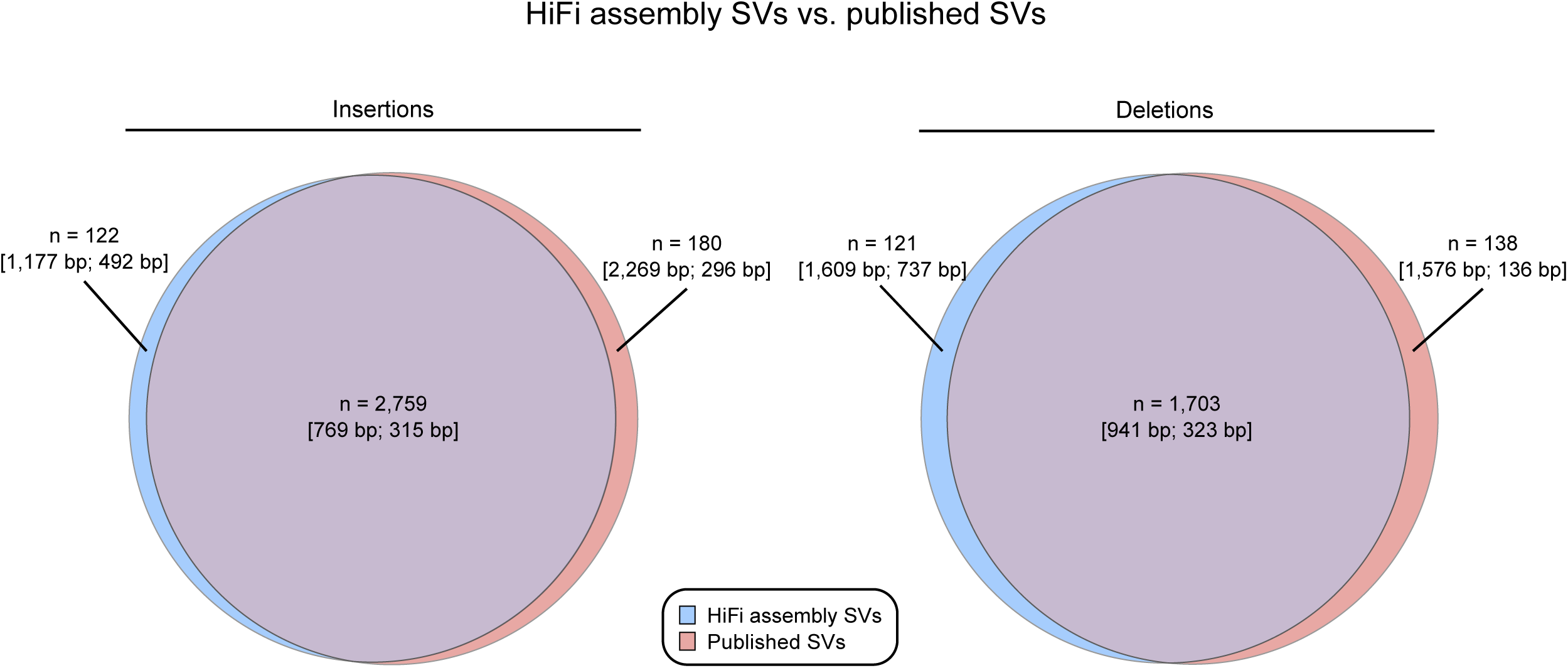
SVs discovered in the HiFi assembly are supported by published CHM13 calls. We intersected SVs with published CHM13 SVs excluding tandem repeat and segmental duplication (SD) loci, where variant comparisons are more challenging. Both insertions (left) and deletions (right) are strongly supported. Each Venn area is annotated with the total number of variants (n) along with the mean and median variant size, respectively, in brackets. For both insertions and deletions, the HiFi assembly calls more variants around 700 bp to 1 kbp, but the published variants have more calls in the 50-100 bp range as well as more larger calls (10+ kbp). These disagreements are reflected in the mean and medians, and they may be due to assembly errors or differences in mapping and assembly methods used in each study.

**Figure S7.**
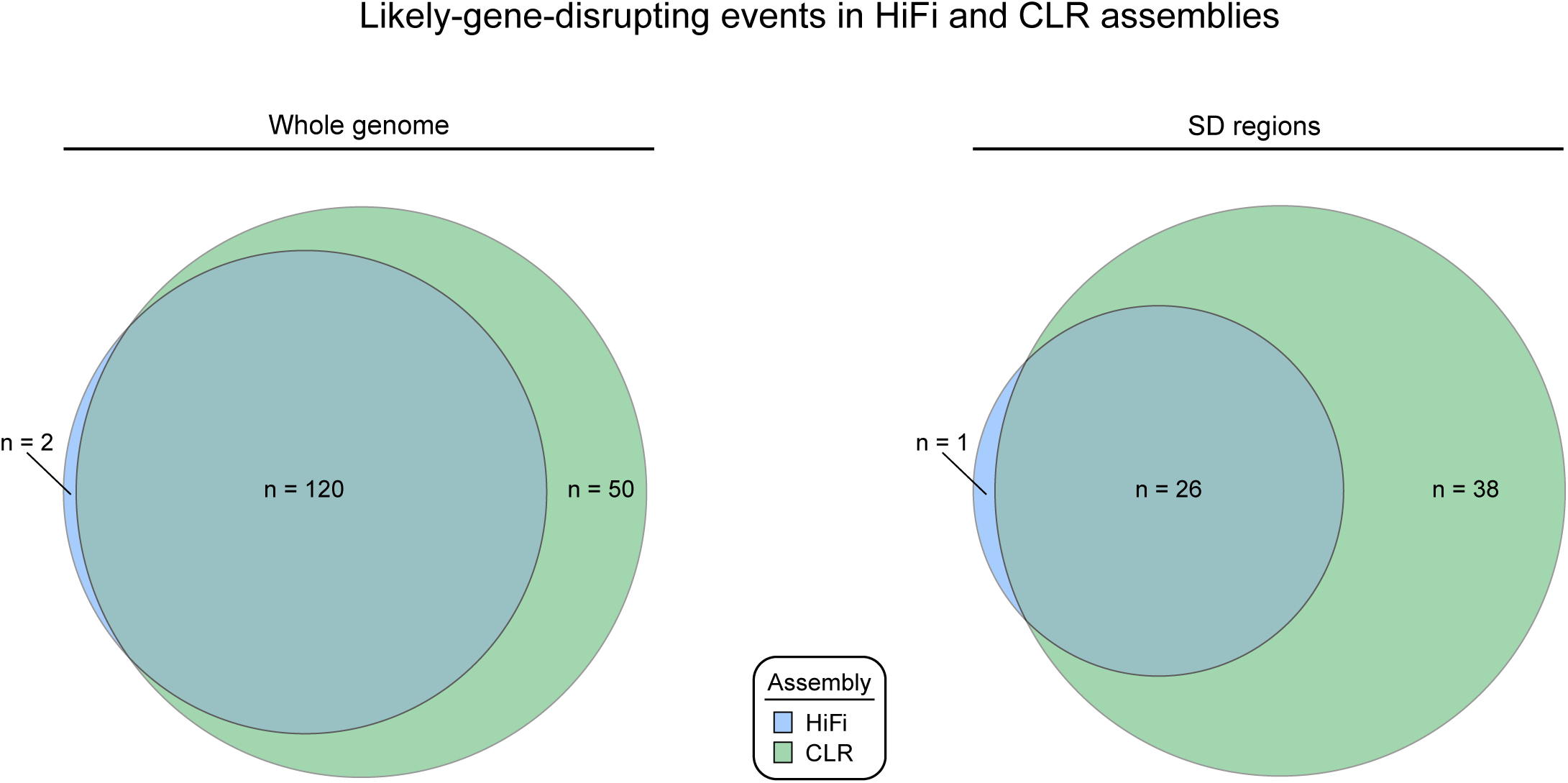
Disrupted genes in the HiFi assembly supported by the CLR assembly. For all loci where the polished HiFi and CLR assemblies had a single alignment to the reference (left), all but two genes disrupted by the HiFi assembly have support in the CLR assembly. We observe 50 genes disrupted in CLR without HiFi support (29% of CLR-disrupted genes). When we restrict our analysis to SDs (right), the percentage of genes disrupted in the CLR assembly without HiFi support increases to 59%.

**Figure S8.**
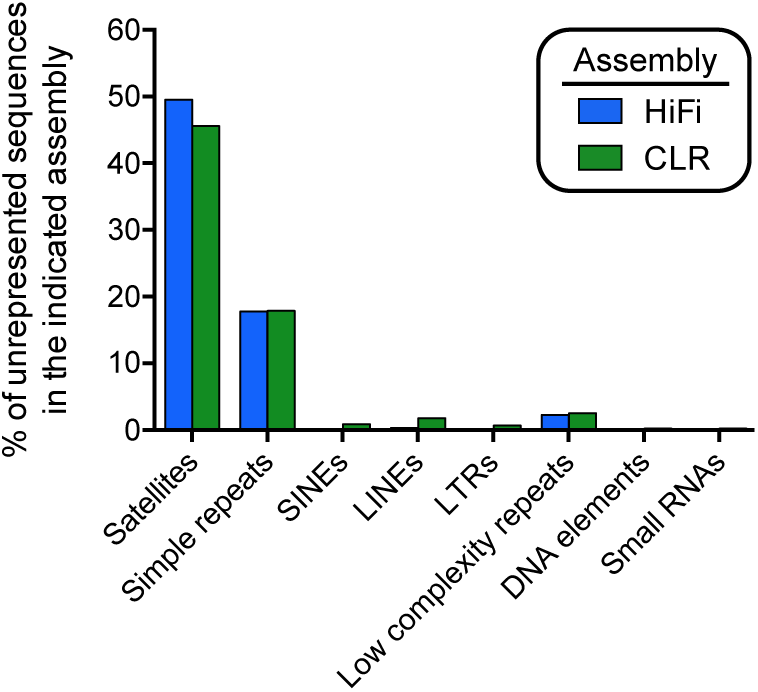
Repeat content of unassembled reads. Bar plot of the repeat composition of sequences not incorporated into the HiFi and CLR assemblies. Most of the unrepresented sequences consist of satellite repeats mapping to heterochromatin or pericentromeric DNA (centromeres, acrocentric DNA and secondary constrictions of chromosomes). SINE, small interspersed nuclear element; LINE, long interspersed nuclear element; LTR, long terminal repeat.

### Supplemental Notes

#### Polishing the assemblies

Initially, the HiFi assembly was polished with Racon using the approximate alignments from minimap2 (i.e., the PAF output generated using the -x asm5 option) and fasta input of HiFi reads. However, we found that this polishing step only modestly increased the QV (by <0.01). When we polished the HiFi assembly with the exact alignments (i.e., the SAM output generated using the -ax map-pb option) and fastq input, we observed a large increase in the median QV (from 40.4 to 45.0). In addition, we observed that the QV achieved using these Racon parameters was greater than that achieved with Arrow (which used >1 TB of CCS subreads). Polishing a second time with Racon further increased the QV and significantly reduced the number of gene-disrupting indels. Adding Pilon polishing did not change the median QV but significantly reduced the total QV across all the BACs because it introduced a 660 bp insertion that appears to be an error relative to the AC275297.1 BAC. Additionally, Pilon polishing only reduced the number of indels genome wide by 645 out of 683,564 (0.094%) and resolved only one additional gene-disrupting event in unique sequence. It is, therefore, our suggestion to polish assemblies generated with HiFi data with two rounds of Racon using the parameters described above rather than with Arrow or Pilon.

#### BAC divergence

In all of our polished assemblies (HiFi or CLR; **Table 1**), we noticed that the same two BACs (AC270121.1 and AC275290.1) had the lowest QV values of those assessed (**Fig. S2**). We examined the alignments of these BACs to all the assemblies and to the HiFi reads and found that these BACs had contractions in tandem repeats relative to the CHM13 cell line. In AC270121.1, there was a 338 bp deletion of a (TCCCCC)n repeat, and in AC275290.1, there was an 80 bp deletion in a (GGCTGAGG)n repeat. In addition, AC270136.1 showed a 62 bp expansion in a poly(T) tract, where the HiFi data supported the HiFi assembly, and AC270122.1 showed an 83 bp insertion, where both Illumina and HiFi data supported the HiFi assembly. Across all 31 BACs used for calculating QV, there was only one mismatched base (AC275285.1:148688-148688). This base appears to be correct in the HiFi assembly, as it was observed in both the HiFi and Illumina data (**Fig. S3**). In combination, these results indicate that many of the BACs with QV < 40 are diverged in sequence when compared to the CHM13 genome due to a mutation that likely arose during BAC generation and/or clonal propagation and do not represent an error in the assemblies. For this reason, our QV values should be interpreted as a lower bound of the true QV.

### Supplemental Tables

**Table S1.** False joins identified by Strand-seq within *de novo* assembly contigs.

| Contig | Start coordinate | End coordinate | No. of false joins |
| --- | --- | --- | --- |
| HiFi |  |  |  |
| tig00000017 | 99,602,681 | 99,618,276 | 1 |
| tig00001347 | 16,388,858 | 16,426,941 | 1 |
| tig00003369 | 22,812,690 | 22,841,943 | 1 |
| tig00002385 | 3,056,424 | 3,204,103 | 1 |
| tig00002385 | 10,092,563 | 10,210,819 | 1 |
| tig00001433 | 2,544,909 | 2,612,082 | 1 |
| tig00004976 | 785,171 | 805,928 | 1 |
| CLR |  |  |  |
| NTIA01000004.1 | 51,797,117 | 51,869,304 | 1 |
| NTIA01000039.1 | 16,068,231 | 16,072,573 | 1 |
| NTIA01000061.1 | 12,087,358 | 12,166,683 | 1 |
| NTIA01000067.1 | 4,639,556 | 4,640,945 | 1 |
| NTIA01000093.1 | 746,665 | 783,795 | 1 |

HiFi: HiFi assembly CLR: CLR assembly

**Table S2.** Comparison of PSVs linked with SDA in HiFi and CLR assemblies.

| Locus | Expected no. of<br>paralogs | Expected no. of<br>phased bases<br>(kbp) | No. of paralogs<br>by SDA<br>(HiFi, CLR) | No. of bases<br>phased by SDA<br>(HiFi, CLR; kbp) | Average length of<br>SDA-phased block<br>(HiFi, CLR; kbp) | No. of PSVs<br>(HiFi, CLR) |
| --- | --- | --- | --- | --- | --- | --- |
| *OPN1LW* | 2 | 70 | 0, 1 | 0, 20 | 0, 20 | 0, 14 |
| *NOTCH2NL* | 5 | 500 | 9, 4 | 443, 345 | 49, 86 | 728, 482 |
| *SRGAP2* | 4 | 520 | 5, 5 | 493, 494 | 99, 99 | 1194, 821 |
| *FCGR2/3* | 3 | 220 | 3, 4 | 141, 140 | 47, 35 | 611, 139 |
| *KANSL1* | 2 | 280 | 4, 3 | 54, 150 | 13, 50 | 48, 92 |

